# Diseases and invasive species have synergistic effects with other anthropogenic threats on the functional and phylogenetic diversity in Testudines and Crocodilia

**DOI:** 10.1101/2024.08.02.606360

**Authors:** R.C. Rodríguez-Caro, R. Gumbs, E. Graciá, S. P. Blomberg, H. Cayuela, M.K. Grace, C.P. Carmona, H.A. Pérez-Mendoza, A. Giménez, K.J. Davis, R. Salguero-Gómez

## Abstract

Understanding how multiple threats interact is crucial for the prioritization of conservation measures. Here, we investigate how interactions between six common threats (climate change, habitat disturbance, global trade, overconsumption, pollution, and emerging diseases/invasive species) affect the functional and phylogenetic diversity of 230 species of Testudines and 21 of Crocodilia. We classify two-way threat interactions into additive, synergistic, and antagonistic according to their effects on functional and phylogenetic diversity. Most threat interactions are antagonistic, the effect of threats jointly is lower than the sum of the effects of threats separately. However, we find that the interaction between emerging diseases or invasive species with other threats has synergistic and additive effects, meaning that the combined effects are greater or equal to the effects of threats separately. Our work can help target conservation strategies and detect key places to address multiple threats when they appear together.

## INTRODUCTION

Habitat loss, climate change, pollution, and overexploitation for consumption and trade are each placing significant pressure on the persistence of species across the globe (Díaz et al., 2019; Maxwell et al., 2016; Steffen et al., 2011). These threats, however, often interact to affect biodiversity, sometimes amplifying their effects and creating additional challenges for species viability (Côté et al., 2016). For example, habitat loss can make species more vulnerable to the impacts of climate change (MantykalJpringle et al., 2012), while poaching and unsustainable trade can further exacerbate the impact of habitat loss in species like the jaguar (*Panthera onca;* RomerolJMuñoz et al., 2019) or elephants (*Loxodonta spp*; Breuer et al., 2016). Despite the impacts of these complex interactions, we currently lack a global understanding of where and which threats interact, and in what direction, to shape functional and phylogenetic diversity. Crucially, this knowledge is critical for the effective implementation of conservation measures (Craig et al., 2017).

Ongoing biodiversity decline is leading to the erosion of functional diversity (Carmona et al., 2021; Toussaint et al., 2021). Reduction in functional diversity, defined in our study as the loss of life history strategies, can reduce the resilience of the ecosystem and may result in the loss of ecological processes (Oliver et al., 2015). In this context, life history strategies describe species’ life cycles and their suitability for adaptation to a given environment (Capdevila et al., 2020; Healy et al., 2019; SalguerolJGómez, R. et al., 2016; Stearns, 1999). We recently examined how the theoretical extinctions of species of Testudines and Crocodilia facing anthropogenic threats would affect the functional diversity and the vulnerability of specific vital life history strategies (Rodríguez-Caro et al., 2023). In that study, we found that different human threats affect specific life histories and therefore differentially risk the functional diversity of the taxonomic group. For instance, species of Testudines and Crocodilia with ‘slow’ life histories (*i.e.*, late maturity and low numbers of offspring), are particularly vulnerable to threats from invasive species and diseases. However, the effects of interactions among multiple threats on functional diversity loss remains unknown.

In addition to functional diversity, assessing losses in phylogenetic diversity can be key to understand evolutionary potential (Faith, 1992) and to prioritize conservation measures (Faith, 2008; Rosauer et al., 2017). Phylogenetic diversity is defined as the total length of the branches on a phylogenetic tree that connect a group of species to each other (Faith, 1992). Phylogenetic diversity assesses the combined influence of species on the overall Tree of Life, measuring the extent of evolutionary variation within a set of species (Faith, 2008). However, there has been little exploration of the expected effects of anthropogenic threats on phylogenetic diversity. Previous work has evaluated the loss of phylogenetic diversity in reptiles relative to the Human Footprint Index (Venter et al., 2016), across the spatial distribution of the species (Gumbs et al., 2020). However, to date no study has specifically explored the effects of interactions of threats on Testudines and Crocodilia species.

Importantly, the outcome of threat interactions can depend on the spatial context of each threat (Capdevila et al., 2022; Bowler et al., 2020). For example, local threats such as habitat destruction or emergent diseases are directly associated with human populations (Di Giulio et al., 2009; Berry et al., 2015). Consequently, the prevalence of these local threats is expected to vary with human population density regardless of latitude (Santini et al., 2017). Globally, however, other global threats like climate change are distributed unevenly, particularly latitudinally (Harfoot et al., 2021; IPCC, 2021). The uneven distribution of certain threats poses challenges in predicting the spatial distribution and the effects of multiple, interacting threats. Moreover, the co- occurrence of threats may endanger the conservation of functional and phylogenetic diversity (Geary et al., 2019).

Testudines (tortoises, terrapins, freshwater and sea turtles) and Crocodilia (crocodiles, alligators, and gharials) have recently been identified as the groups whose functional and phylogenetic diversity are most threatened by extinction (Rhodin et al., 2018; Colston et al., 2020; Gumbs et al., 2020; Rodríguez-Caro et al., 2023). Indeed, 50-60% of species in both groups are threatened with extinction (Cox et al., 2022; IUCN, 2020). Moreover, the extinction of these species could result in greater-than- expected losses in functional diversity (Rodríguez-Caro et al., 2023). To make things worse, the interactions among multiple anthropogenic threats to these threatened species are exceptionally high. For example, the Roti Island snake-necked turtle (*Chelodina mccordi*), a Critically Endangered species (As-singkily et al., 2019), is threatened by habitat disturbances, which have resulted in its displacement towards more anthropogenic areas where the risk of illegal trade for pet collection is higher (Rhodin et al., 2018). These threats have been compounded by the emergence of exotic species, pollution, and climate change, which pose a significant risk to the persistence of this (Rhodin et al., 2004; Eisemberg et al. 2016) and other species (Cox et al., 2022).

Here, we quantify how interactions between different threats may alter the functional and phylogenetic diversity of Testudines and Crocodilia, and identify the regions around the world with higher risk of said loss. To do so, we first analyse the loss of functional and phylogenetic diversity by simulating the extinction of species affected by various anthropogenic threats and quantifying the associated loss of functional and phylogenetic diversity. Next, according to Côté et al. (2016), we assess whether the effects of pair-wise combinations between different threats exhibit different relationships: (i) Additive: the loss of functional or phylogenetic diversity from each threat separately is equivalent to the loss of diversity when both threats act simultaneously. An additive effect would indicate that the threats affect different domains of the functional spectra or disparate evolutionary lineages (Fig. 1); (ii) Antagonistic: the loss of functional or phylogenetic diversity of two threats together is lower than the threats separately. An antagonistic effect would indicate that similar species are affected by the same threats (Fig. 1); and iii) Synergistic: the combined loss of functional or phylogenetic diversity when the two threats act jointly is greater than when two threats affect separately. A synergistic effect would indicate that the threats affect complementary species in the same region of the functional space or phylogenetic tree, completely eliminating this part of the space (Fig 1). We hypothesise that, in general, (H1) the loss of functional and phylogenetic diversity resulting from multiple threats exhibits antagonistic effects. However, we expect that (H2) interactions among threats that are known to affect specific portions of the functional spectrum of testudines and crocodilians, such as local consumption, diseases and invasive species, and pollution (Rodríguez-Caro et al., 2023), may exhibit additive or synergistic effects.

**Fig. 1.**
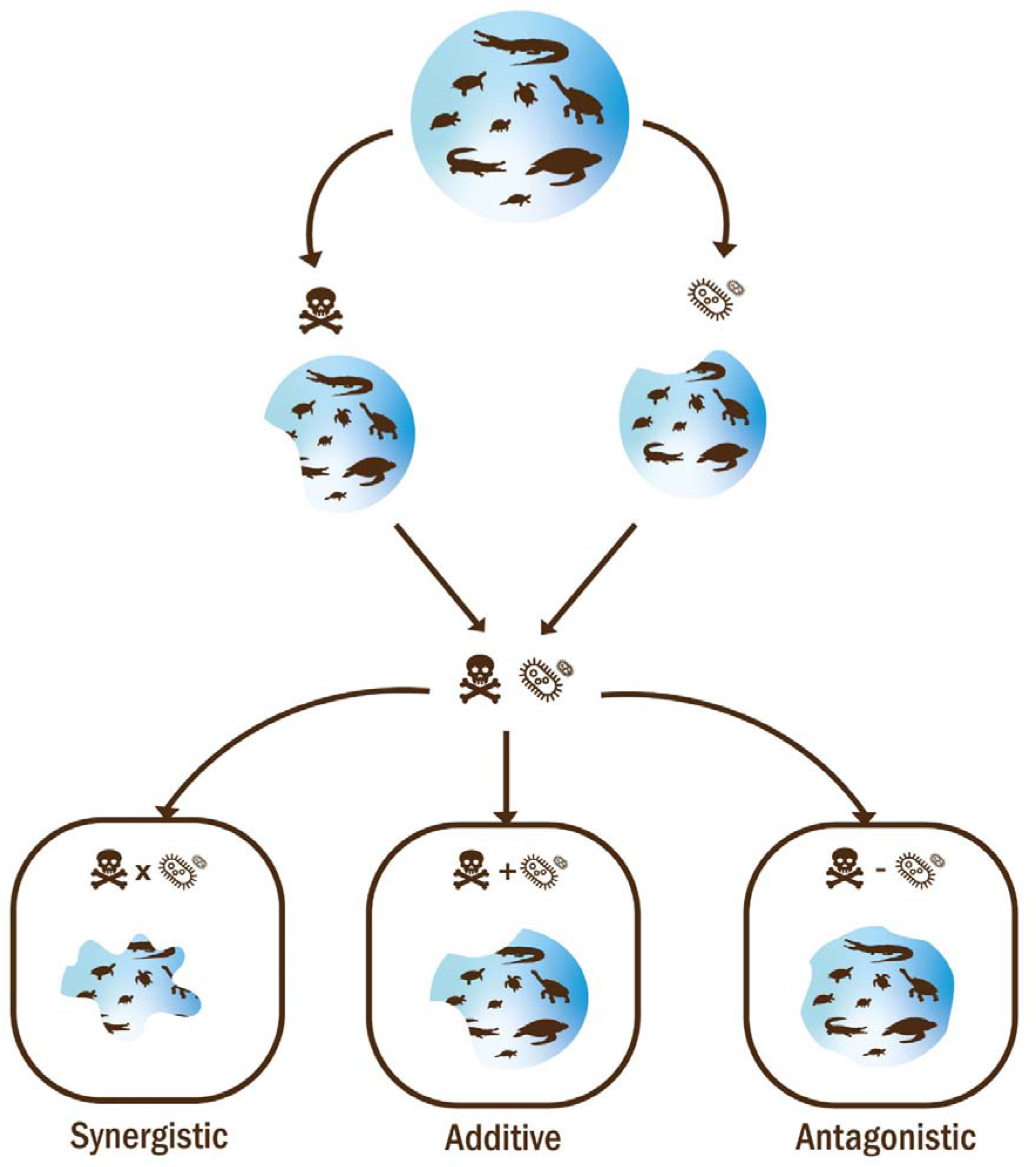
Diagram illustrating the three types of threat interactions observed in our analyses of loss of functional and phylogenetic diversity. The effects of combinations pairwise threats can in principle exhibit three different outcomes: (1) Synergistic: the combined loss of functional or phylogenetic diversity when two threats act jointly is higher than the loss diversity of the two threats separately; (2) Additive: the loss of functional or phylogenetic diversity from each threat separately is equivalent to the loss of diversity when both threats act simultaneously; and (3) Antagonistic: the combined effect when both threats act together is lower than the loss of functional or phylogenetic diversity of two threats analysed separately.

## MATERIALS AND METHODS

### Life history strategies, functional spectra, and phylogenetic diversity

To identify the functional trait space and examine functional diversity of Testudines and Crocodilia, we quantified their life history strategies using life history trait data. Life history traits define key moments along the life cycle of a species (*e.g.*, age at maturity, maximum longevity; Capdevila & Salguero-Gomez 2021), and are underpinned by the vital rates of survival, development, and reproduction (Roff et al., 1993). We obtained these trait data from the published literature, as detailed in Rodríguez-Caro et al. (2023).

Briefly, life history traits were obtained from COMADRE Animal Matrix Database v. 4.23.3.1 (Salguero-Gómez et al., 2016), DATLife Database (DATLife Database, 2021), Amniote Life History Database (Myhrvold et al., 2015), and published reviews (Allen et al., 2017; Pfaller et la., 2018; Reinke et al., 2022). Missing traits in the dataset (38%, Fig. S1) were imputed using the R package *mice* (Van Buuren & Groothuis-Oudshoorn, 2011), which uses multiple imputation, and the add-on *phylomice* to include phylogenetic information (Rodriguez-Caro et al., 2023). We used six life history traits that encompass detailed information regarding the timing, intensity, frequency, and duration of vital rates across the life cycle of any species: adult survival (*Sa*), juvenile survival (*Sj*), maximum lifespan (*ML*), age at sexual maturity (*L*α), mean number of clutches per year (*CN*), and clutch size (*CS*). The resulting data encompass 259 species: 236 testudines and 23 crocodilians.

To identify potential differences between the patterns of association among life history traits for Testudines and Crocodilia species, we used a phylogenetically informed PCA (*p*PCA), corrected by adult body mass. This approach allows us to examine functional diversity while also assessing and estimating the strength of non- independence among the examined lineages (Revell, 2009). The *p*PCA considers the correlation matrix of species’ traits while accounting for phylogenetic relationships and simultaneously estimates phylogenetic signal via Pagel’s λ (Freckleton, 2000), which ranges from 0 (not explained by phylogeny) to 1 (the observed pattern is correlated with the phylogeny). The *p*PCA was estimated using the R package *phytools* (Revell, 2012), assuming a Brownian motion model of evolution (Revell, 2010). Life history trait data were loglJtransformed to fulfil normality assumptions of PCA and zlJtransformed to meanlJ=lJ0, and SDlJ=lJ1 (Legendre and Legendre, 2012). To account for the potential effect of body mass in life history analyses (data from Rodríguez-Caro et al., 2023), we used the residuals from a phylogenetic regression of adult body size and each trait.

To describe the functional spectra of our study species, we estimated the multivariate kernel density of the trait data. To do so, we used the *TPD* (Trait probability density; Carmona et al., 2019) and *ks* R packages (Duong, 2007, 2014) for the two first axes of the *p*PCA (Rodríguez-Caro et al., 2023). The resulting grouped kernels for all species were transformed into the continuous TPD function. Following methods described in detail elsewhere (Carmona et al. 2021; Toussaint et al., 2021 and Rodríguez-Caro et al., 2023), we divided the continuous functional space into a two- dimensional grid composed of 200 equal sized cells per dimension. Next, we estimated the value of the TPD function for the 40,000 cells. In this way, the value of the TPD function represents the density of species in that particular region of the functional space (*i.e.*, species with similar life history strategies).

We used the phylogenetic trees of Colston et al. (2020) to represent the phylogenetic relationships of the species in our study. We sampled 1,000 phylogenetic trees from the published set of 10,000 to adequately reflect the variation in phylogenetic placements and divergence time estimates inherent in such ‘pseudo-posterior’ distributions of trees (Thomas et al. 2013). The sum of all branch lengths of the phylogenetic tree represents the total PD of the clade (Faith 1992).

### Threats to the studied species

To identify threats of Testudines and Crocodilia species, we used data carefully collated from three sources, detailed in Rodriguez-Caro et al. (2023). Briefly, these sources are Stanford et al. (2020), Bonin et al. (2006) and the section about threats in the IUCN Red List (2020). The main described threats were: (i) habitat loss, fragmentation, and degradation (Luiselli, 2009), (ii) over-collection of individuals and their eggs for food consumption (Gong et al., 2017), (iii) unsustainable or illegal international trade, as well as over-collection for the trade in medicines (Sung and Fong, 2018), (iv) climate change (Gibbons et al., 2000); (v) invasive species and emergent diseases (Jacobson et al., 2014; Tompkins et al., 2015); and (vi) pollution (Hutchinson and Simmonds, 1991). We assessed these six threats for 230 (65% of the extant species) species of Testudines, and for 21 (78% of extant species) species of Crocodilia. Due to the lack of data regarding the intensity of threats to each species, here threat variables were categorised as 1 (or 0) if a specific threat was described for the species (or not).

### Effects of interactive threats on global functional and phylogenetic diversity

To quantify the effect of the threats on the functional spectra of our 251 species of Testudines and Crocodilia, we simulated threat-specific extinction scenarios. In each scenario, species were classified as extinct if they were reported as affected by the assessed threat. To evaluate the differences between threatened and non-threatened species, we carried out two comparisons: (1) between the TPD function considering all the species (current spectra of functional diversity) and the TPD function after removing the species affected by each of the six threats separately (habitat degradation, trade, local consumption, climate change, or interactions with invasive species and diseases);, irrespective of their threatened category; and (2) between the TPD function considering all the Red List-assessed species, and the TPD function after removing the threatened species (Critically Endangered (CR), Endangered (EN), or Vulnerable (VU)) affected by each specific threat. Next, we estimated the loss of functional spectra using the TPD function considering all the species assessed, and the TPD function after removing the species according to specific threats. Here, the comparisons between TPD functions are supported by the fact that they are probability density functions, and as such they integrate to 1 across the whole functional space, regardless of the number of species considered (Carmona et al., 2016). To reduce the potential effect of outliers in the functional space, we applied a quantile threshold of 99%, following Carmona et al. (2016). Finally, we quantified how much of the functional spectra is lost after the extinction scenarios according to the different threats by estimating which functional space cells became empty after our simulated extinctions.

To identify the type of relationship between the different threats (hypotheses H1 and H2 outlined in our introduction), we estimated the potential loss of functional diversity attributable to each threat and between the pairwise combinations of threats. To that end, we use a comparison between TPD functions similar to the one explained above. We then evaluated the differences among single and multiple effects to identify synergic, antagonistic, and additive effects (Figure 1). We carried out these comparisons both considering the extinction of all the species affected by each threat and considering only the extinction of threatened species (CR, EN, VU). To provide a spatial description of the interactions between threats, once the relationships between the different threats (antagonistic, synergistic, or additive) have been described, we identified on which continents these relationships occur at the species level. We estimated how often synergists, antagonistic, and additive effects appear in each continent to assess which ones face higher risks of functional diversity loss in Testudines and Crocodilia species.

We repeated the same scenarios of extinction based on single and pairwise combinations of threats across the 1,000 phylogenetic trees. For each phylogenetic tree, the total phylogenetic diversity (PD) was calculated prior to the removal of the species affected by each threat or combination of threats. The phylogenetic diversity was then recalculated following the removal of the affected species from the phylogeny and the difference in phylogenetic diversity values (*i.e*., the amount of phylogenetic diversity lost) was calculated for each scenario across each of the 1,000 trees.

## RESULTS

### Effects of interactions

To assess the differences between the individual effects of threats and their combined effects, we compared the loss of functional diversity in each case. By conducting pairwise comparisons among the different threats, we determine that most combinations of threats exhibit antagonistic effects (73.3% of combinations, Fig. 2). However, one pairwise combination of threats shows an additive effect (pollution × invasive species/diseases), and 20% of combinations exhibit synergistic effects, such as invasive species/diseases × habitat disturbances, invasive species/diseases × local over- consumption, and invasive species/diseases × international trade. Additionally, while the analysis of the relation between threats remains consistent between the two subsets of data (threatened and total species), differences were found in the combination of unsustainable local consumption × habitat disturbance. In that combination, the relationship is additive in the analyses with all species, but when considering only the threatened species (CR, EN, and VU), the relationship becomes antagonistic (Fig 2a).

**Fig. 2.**
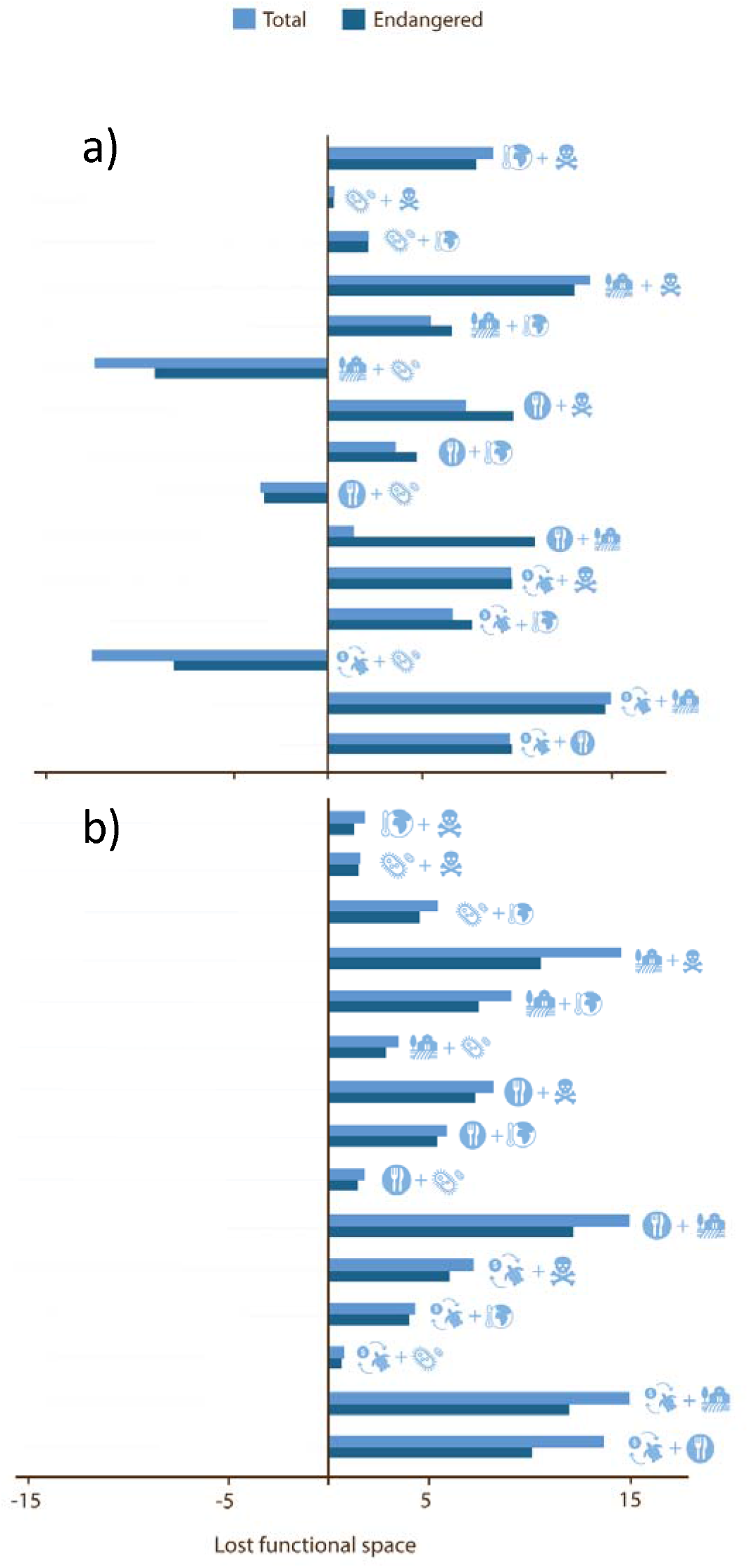
Most of the threat interactions that shape the loss of functional and phylogenetic diversity in Testudines and Crocodilia worldwide are antagonistic. Positives values of loss of functional space is related to antagonistic relationships, negative values are related to synergistic interactions and values near to zero represent additive interactions. However, synergistic or additive relationships are observed when diseases and invasive species are involved for a) functional diversity and b) phylogenetic diversity. A) The bar chart illustrates the effects on functional diversity loss by comparing the individual effects of each threat with the combined effects of all threats acting simultaneously. B) Bar chart with the loss of phylogenetic diversity with the combined effects of threats. Negative values indicate synergistic effects, positive values indicate antagonistic effects, and values close to 0 indicate additive effects. The analysis has been performed twice, once with the entire species set (light blue) and once with the threatened species subset (dark blue).

When examining the effect of pairwise interactions of threats on phylogenetic diversity, practically all relationships are antagonistic (Fig 2b). This means that the combined effect of both threats on phylogenetic diversity is lower than the effect of threats separately (results about the loss of phylogenetic diversity separately are in Supplementary materials). However, when the interaction occurs between the threat of emerging diseases and invasive species with unsustainable global trade, the loss of phylogenetic diversity shows additive results. Hence, different threats affect divergent evolutionary lineages.

### Spatial evaluation of interactions

To evaluate the spatial distribution of the different relationships among threats, each pairwise combination has been classified into three categories (additive, synergistic, and antagonistic). In the spatial results, in most regions, the highest proportion of effects is antagonistic, indicating that the same species are affected by multiple threats. However, synergistic effects are more prominent in species from Oceania, accounting for 20% of the interactions, as well as in marine species distributed worldwide, representing 16% of the interactions. Despite additive effects being observed in only one type of interaction, their representation is as much as 10% of the interactions in North America (Fig 3).

**Fig. 3.**
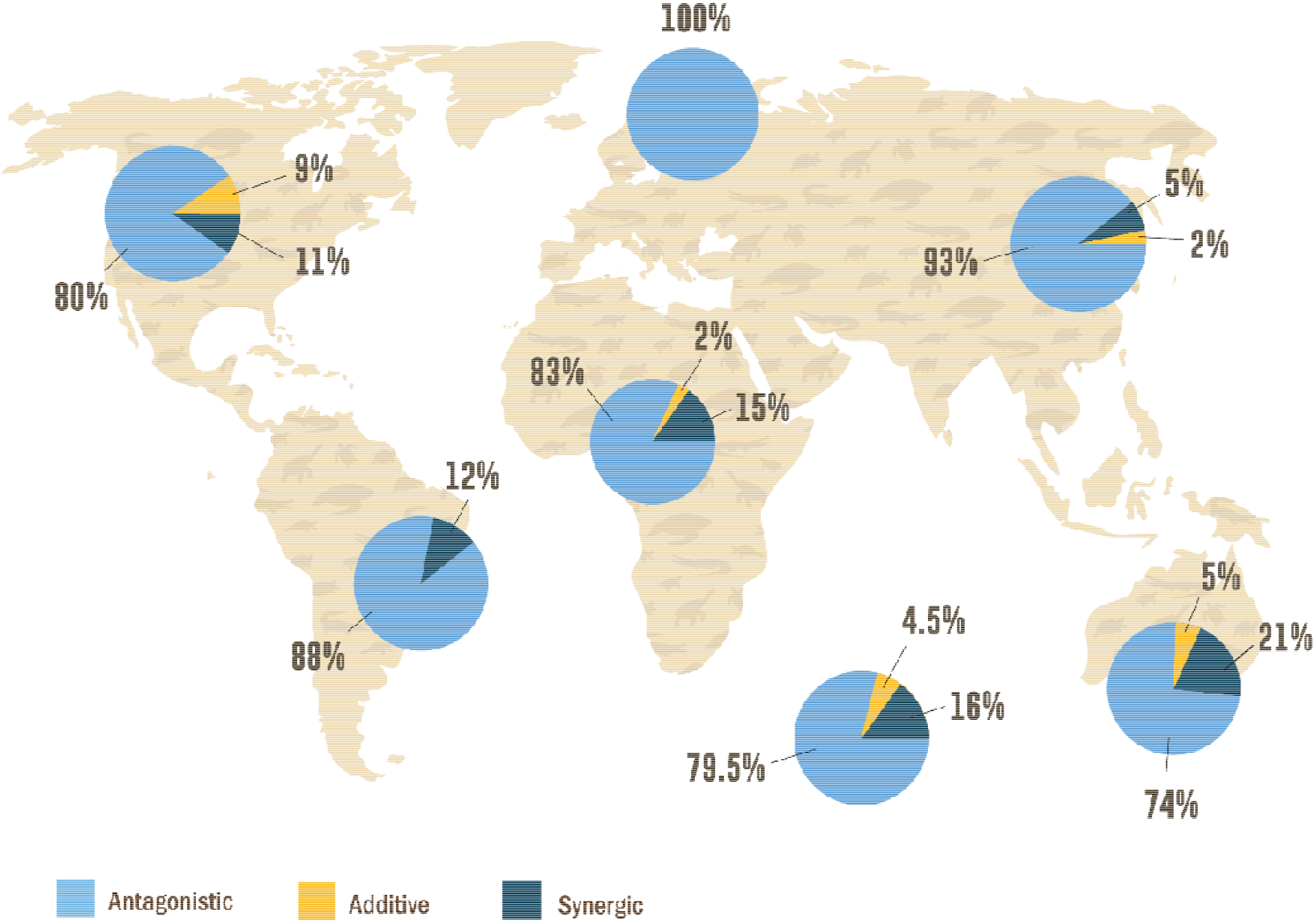
Despite the high occurrence of antagonistic effects of human threats on the functional diversity of Testudines and Crocodilia worldwide, synergistic effects are quite common in Oceania. The pie chart illustrates the proportion of each interactive pair-wise threat effect in different continents. The threats (not shown) are habitat degradation, unsustainable consumption, illegal trade, climate change, interaction with invasive species and diseases, and pollution. Bottom centre pie chart represents worldwide distributed marine species.

The additive effects found in the loss of phylogenetic diversity occur mainly in two regions, North America and Africa. In these regions, the threats of emerging diseases and invasive species with unsustainable trade have a higher occurrence (42% occur in North America and 33% are Africa), with the remaining continents showing lower representation.

## DISCUSSION

Here, we analysed the interaction of multiple threats on the functional and phylogenetic diversity of two groups of high conservation interest: Testudines and Crocodilia (Colston et al., 2021). We report a wide disparity in the effect of threats on both groups, with prevalent antagonistic interactions between threats (73.3% of threat combinations), whereby the impact of two threats together is lower than the reduction in diversity when these threats are considered individually. Additionally, we found differential distributions of the threat interactions across the globe, with a high proportion of synergistic interactions in Oceania. Finally, our analyses show that variability in said interacting effects is greater concerning functional diversity than phylogenetic diversity.

The fact that most threats present antagonistic relationships (hypothesis H1) reflects the high redundancy in life histories in reptiles (Carmona et al. 2021, Rodríguez-Caro et al., 2023). Our results indicate that the same species is indeed affected by several threats simultaneously. For instance, in the case of the genus *Testudo*, species are mainly threatened by habitat loss, climate change, and illegal trade (Graciá et al., 2020). In fact, in our results, 43% of our examined species for which threats have been described are at risk from at least three different threats. In our database, species such as the Olive Ridley sea turtle (*Lepidochelys olivacea*) and The Roti Island snake-necked turtle (*Chelodina mccordi*) face risks from all six threats examined in this study. Both species are long-lived threatened species, and the population trend is decreasing according to the Red List (IUCN, 2020). However, the occurrence of additive or synergistic relationships in our study indicates that threats affect complementary functional groups, which points to greater risks of extinction and loss of diversity (Côté et al., 2016). Synergistic relationships described for other taxonomic groups such as neotropical primates, affected by habitat fragmentation and hunting, have helped focus attention on holistic conservation efforts; for example, restoring connectivity can indirectly reduce hunting impacts if human access is restricted to local communities (Mancini et al., 2023). In our case, conservation policies aiming to preserve testudines and crocodilians should be directed not towards specific species, but towards key threats to the conservation of functional diversity. For example, the management of invasive species must accompany habitat restoration, as we have found a synergistic relationship among these threats.

In our study, synergistic relationships usually are found when one of the threats is invasive species/emerging diseases, thus providing support for our hypothesis H2. Emerging diseases and invasive species affect slow testudines and crocodilians (*i.e.* species with long generation length and maximum lifespans such as terrestrial tortoises) more than fast ones (Rodríguez-Caro et al., 2023). However, when Testudines or Crocodilia species are also affected by other more generalist threats, such as habitat loss or unsustainable trade, the loss in functional diversity is much greater than expected (Cox et al., 2022; Rodríguez-Caro et al., 2023). Here, emerging diseases, such as diseases in the respiratory tract in tortoises (Origgi and Jacobson, 2020), interact synergistically with other globally-distributed threats like habitat disturbances (Farooq et al., 2023), and together they may pose a high risk to the loss of functional diversity of this taxonomic group.

We found that synergistic interactions of threats occur mainly in Australia and in worldwide seas. In Australia, invasive species are widely recognised as the second largest threat to the local biodiversity (Evans et al., 2011). Australia is one of the most important hotspots of biodiversity in the world (Lindenmayer et al., 2010), but also has one of the highest rates of species extinction (Woinarski et al., 2015). Previous studies have described that the co-occurrence of threats to biodiversity in Australia jeopardizes species conservation (Allek et al., 2018). In fact, Australian vertebrates face an average of six threats, with habitat fragmentation and invasive species being the most common ones (Hobbs, 2002; Allek et al., 2018). On the other hand, our results show that marine species are also affected by synergistic effects of threats, for sea turtles, the first life stages are particularly sensitive to invasive species (Stokes et al., 2024), habitat disturbances (Mathenge et al., 2012), and local overconsumption in beaches with human presence (López-Mendilaharsu et al., 2020). Indeed, recent studies have highlighted the importance of considering that threats in marine and terrestrial ecosystems may interact synergistically such as with loggerhead sea turtles (*Caretta caretta)* in the Mediterranean basin (Mancino, et al., 2022). Therefore, establishing measures to control this combination of threats in regions like Australia or in worldwide seas is urgent to halt the loss of functional diversity.

Although the loss of functional diversity shows synergistic or additive interaction, the loss of functional diversity by multiple threats is mainly antagonistic for Testudines and Crocodilia. Given the hierarchical nature of phylogenetic trees, a large proportion of the phylogenetic diversity of a clade is typically contained along internal branches ancestral to multiple species (Faith, 2008). Previous studies have found that phylogenetic diversity correlates with functional diversity when threats are examined in isolation in plants (Tucker et al., 2018), and in mammals (Brodie et al., 2021). These findings support our results for Testudines and Crocodilians because both diversities show similar correlations (Rodríguez-Caro et al., 2021; Supplementary material 1). However, here we show that phylogenetic diversity is affected by multiple threats in additive and antagonistic ways, whereas functional diversity also shows synergistic effects. For synergistic effects to emerge from two threats, all species in a given clade must be impacted by at least one of them threats: a single species unaffected by either threat will prevent the loss of all branches from which it descends. Therefore, synergistic effects on phylogenetic diversity are expected to be rare. However, the emergence of synergistic effects would represent a serious risk of tipping points leading to the loss of deep phylogenetic branches (Faith 2015).

Analysing the impact of interactions between threats on species viability is crucial not only to prioritise conservation measures, but also to detect hotspots where conservation efforts are urgently needed (Craig et al., 2017). Here, we analysed threats pairwise to aid in tractability. However, complex models with larger combinations of threats could help identify specific life history strategies at risk, as well as locations in the world where various threats may co-occur. Our analyses are limited to threatened species for which some threats have been described, but our approach allows us to identify which life history strategies may be affected and extrapolate their effects to other species with less information available, for example, using new databases about the information of threats worldwide in reptiles (such as Farooq et al., 2023). Expanding our approach to other taxonomic groups may be crucial for rapid threat detection and to improve conservation of threatened species for which threats have not yet been described. This approach will be of special interest for reptiles and amphibians, for which significant gaps in information exist (Conde et al., 2019). Our study serves as a starting point for assessing the effects of interactions between threats on wildlife.

## ACKNOWLEDGEMENTS

RCRC was supported by the European Union-Next Generation EU in the Maria Zambrano Program (ZAMBRANO 21-26). RSG was supported by a NERC Pushing the Frontiers grant (NE/X013766/1). CPC was supported by the Estonian Research Council (PRG2142 and MOBERC100) and the European Union (ERC, PLECTRUM, 101126117). EG and AG was supported by Project TED2021-130381B-I00, funded by the Spanish Ministry of Innovation, Science and Universities (MICIU/AEI/10.13039/501100011033), with the support of the European Union “NextGenerationEU”/PRTR”. Illustrations by Carmen Cañizares (@canitailustradora).

## AUTHOR CONTRIBUTIONS

R.C.R.-C. and R.S.-G. conceived the original idea. R.C.R.-C. curated the data. R.C.R.-C. and R.G. R.S.-G. supervised the analyses. R.C.R.-C. and R.S.-G. wrote the first draft and all authors (R.C.R.-C., R.G., E.G., S.P.B., H.C., M.G., C.P.C., H.A.P.-M., A.G., K.D. and R.S.-G.) reviewed and edited the manuscript.

