## Supplementary material for "Diseases and invasive species have synergistic effects with other anthropogenic threats on the functional and phylogenetic diversity in Testudines and Crocodilia"

**Supplementary materials**

### The interaction of emerging diseases and invasive species with other threats has synergistic and additive effects that threat functional and phylogenetic diversity in Testudines and Crocodilia

Rodríguez-Caro, R.C., Gumbs, R., Graciá, E., Blomberg, S. P., Cayuela, H., Grace, M.K., Carmona, C.P., Pérez-Mendoza, H.A., Giménez, A, Davis, K.J., Salguero-Gómez, R.

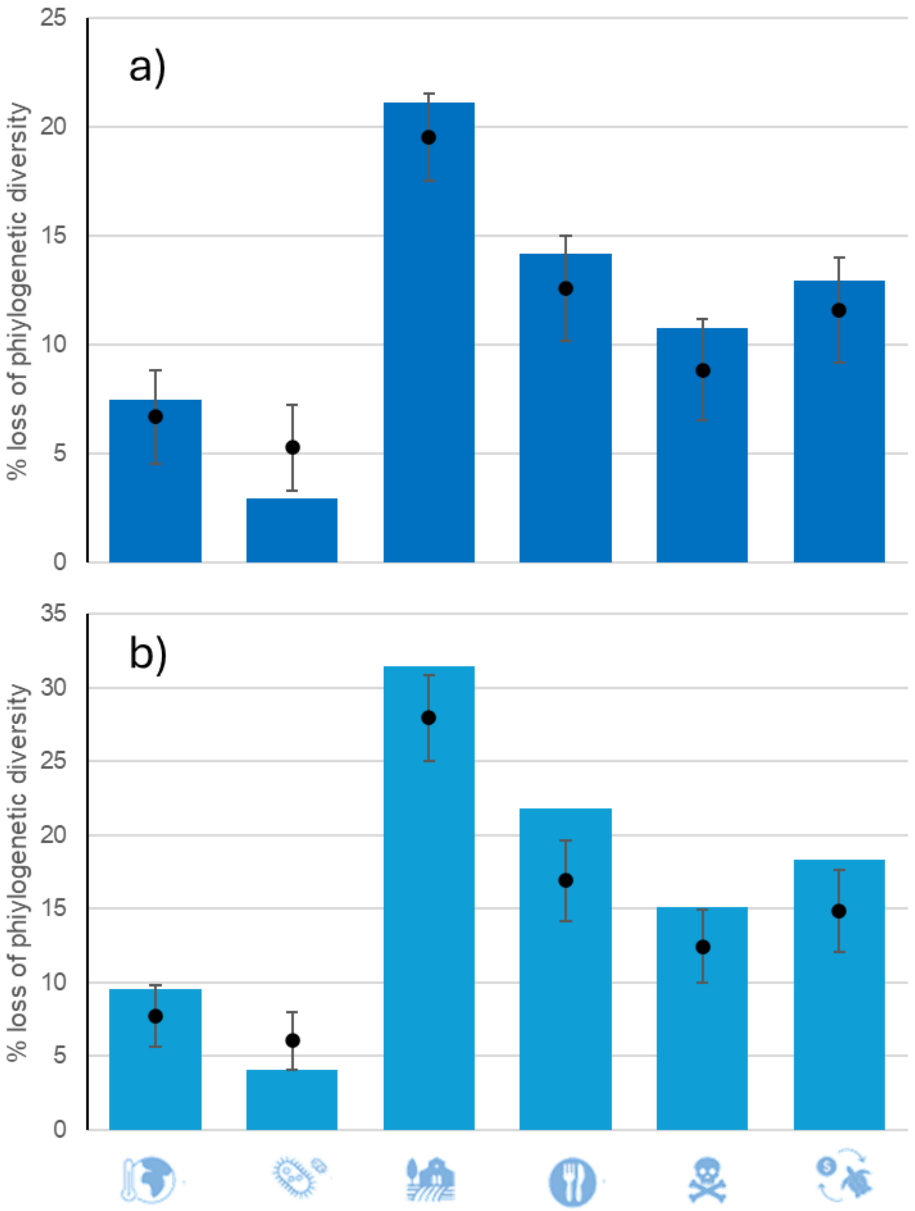


**Fig 1S. Loss of phylogenetic diversity according to the anthropogenic threats.** Simulated loss of phylogenetic diversity of Testudines and Crocodilia under extinction scenarios by threat (N = 251). The loss of functional diversity is expressed as a percentage of the total spectra of functional diversity of species. We simulate the loss of phylogenetic diversity by removing all species affected by specific threats (climate change, diseases/alien species, habitat disturbances, unsustainable consumption, pollution and illegal trade), in a) removing only threatened species (i.e. Critically Endangered [CR], Endangered [EN] and Vulnerable [VU] as per the IUCN Red List) and in b) removing all the species affected (threatened or not). For each scenario, we compare the loss of phylogenetic diversity with 999 iterations of a null model where the same number of species were randomly selected among all 251 species. The 999 randomisations are represented for each threat as a grey dot for the 50th percentile, with grey whiskers representing the 5th and 95th percentiles.

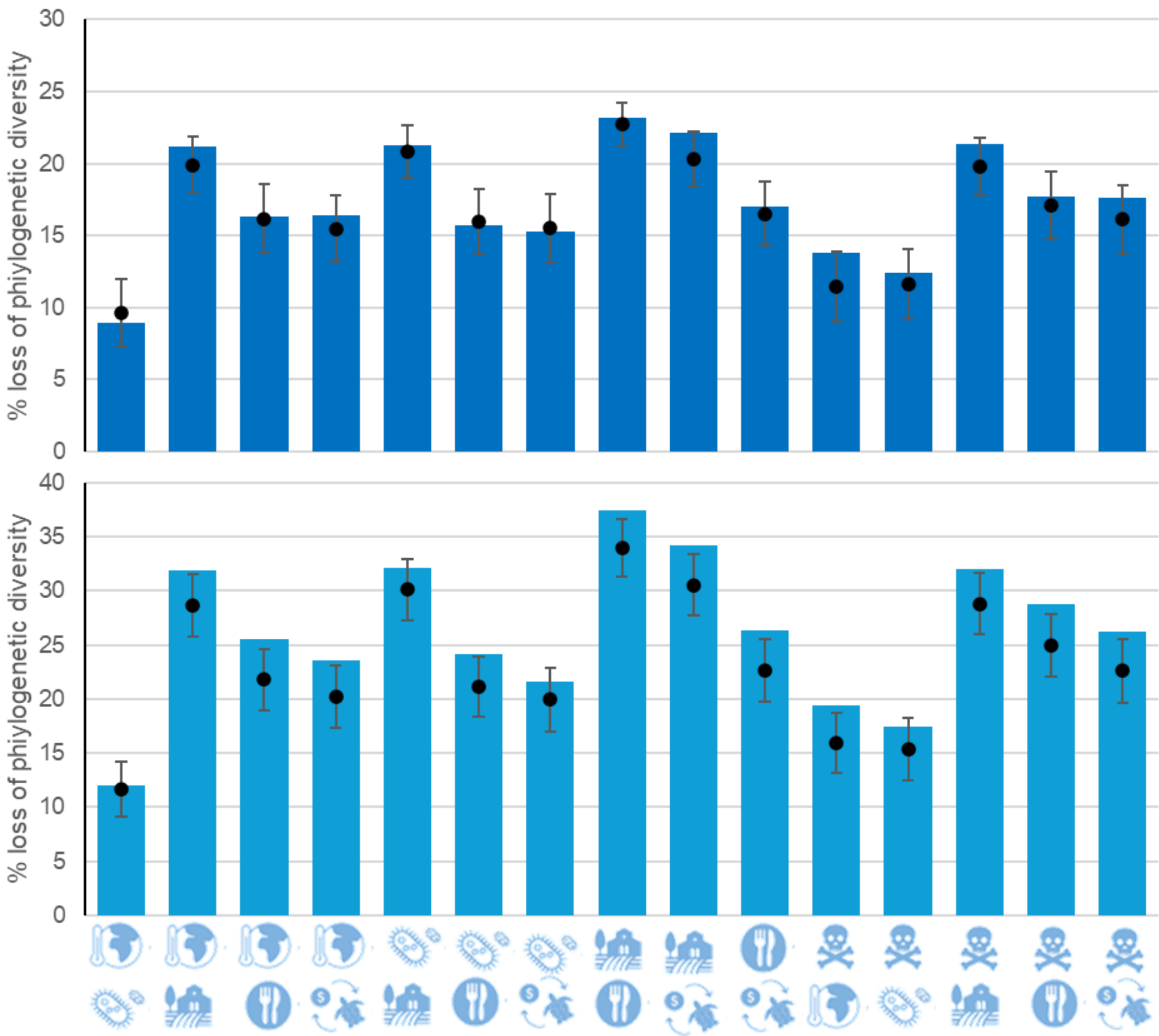


**Fig 2S. Loss of phylogenetic diversity according to the interaction of anthropogenic threats.** Bar chart with the loss of phylogenetic diversity with the combined effects of threats. Simulated loss of phylogenetic diversity of Testudines and Crocodilia under extinction scenarios by the interaction of threats. We a) removed only threatened species (i.e. Critically Endangered [CR], Endangered [EN] and Vulnerable [VU] as per the IUCN Red List) and, b) removed all the species affected (threatened or not). For each scenario, we compared the loss of phylogenetic diversity with 999 iterations of a null model where the same number of species were randomly selected among all 251 species. The 999 randomisations are represented for each threat as a grey dot for the 50th percentile, with grey whiskers representing the 5th and 95th percentiles.
